# Heat stress disrupts acid-base homeostasis independent of symbiosis in the model cnidarian *Exaiptasia diaphana*

**DOI:** 10.1101/2023.05.31.543134

**Authors:** Luella Allen-Waller, Katelyn G. Jones, Marcelina P. Martynek, Kristen T. Brown, Katie L. Barott

## Abstract

Heat stress threatens the survival of symbiotic cnidarians by causing their photosymbiosis to break down in a process known as bleaching. The direct effects of temperature on cnidarian host physiology remain difficult to describe because heat stress depresses symbiont performance, leading to host stress and starvation. The symbiotic sea anemone *Exaiptasia diaphana* provides an opportune system in which to disentangle direct vs. indirect effects of heat stress on the host, since it can survive indefinitely without symbionts. Here, we tested the hypothesis that heat stress directly influences cnidarian physiology by comparing symbiotic and aposymbiotic individuals of a clonal strain of *E. diaphana*. We exposed anemones to a range of temperatures (ambient, +2°C, +4°C, +6°C) for 15-18 days, then measured their symbiont population densities, autotrophic carbon assimilation and translocation, photosynthesis, respiration, and host intracellular pH (pH_i_). Anemones with initially high symbiont densities experienced dose-dependent symbiont loss with increasing temperature, resulting in a corresponding decline in host photosynthate accumulation. In contrast, anemones with low initial symbiont densities did not lose symbionts or assimilate less photosynthate as temperature increased, similar to the response of aposymbiotic anemones. Interestingly, pH_i_ decreased in anemones at higher temperatures regardless of symbiont presence, cell density, or photosynthate translocation, indicating that heat stress disrupts cnidarian acid-base homeostasis independent of symbiosis dysfunction, and that acid-base regulation may be a critical point of vulnerability for hosts of this vital mutualism.

**Summary Statement:** Warming oceans threaten marine invertebrates. We found that heat disrupts acid-base homeostasis in a model symbiotic sea anemone regardless of symbiont presence or function, highlighting bleaching-independent effects of climate change.

## INTRODUCTION

The world’s oceans have absorbed 93% of excess planetary heat from anthropogenic climate change (Johnson and Lyman, 2020), threatening the survival of sessile marine organisms that cannot quickly adapt or migrate to cooler climates to avoid increasingly frequent and severe marine heatwaves (Atkins and Travis, 2010; Kim et al., 2023; Smith et al., 2023). Marine heatwaves have already decimated populations of keystone invertebrates, including notable losses of reef-building corals (scleractinians) (Hughes et al., 2017; Hughes et al., 2018). Predicting how climate change will affect marine invertebrate populations is complicated by the fact that many species, including reef-building coral, engage in symbiotic relationships with microbial symbionts (e.g. dinoflagellates, fungi, bacteria), whose effect on host environmental response we are only beginning to understand (Apprill, 2020; McFall-Ngai et al., 2013). Symbiosis can facilitate physiological adaptation, as partner switching or ‘symbiont shuffling’ can provide mechanisms besides genetic evolution by which organisms can adapt to changing surroundings (Cunning et al., 2015b; Toby Kiers et al., 2010). Yet the interdependence between partners can also magnify the mutualistic organisms’ environmental sensitivities, as both species must maintain function together under new conditions (Apprill, 2020; Bénard et al., 2020; Goulet and Goulet, 2021). We therefore need to be precise in our understanding of when an abiotic stressor like elevated temperature is affecting a host, its symbiont(s), both partners, or even modifying the dynamics of the interaction itself.

Direct effects of abiotic stress on many symbiotic invertebrates remain uncharacterized because environmental change can also have indirect consequences by disturbing obligate symbiosis function. For example, endosymbiotic dinoflagellates (family Symbiodiniaceae) can meet up to 90% of coral energy requirements by providing glucose and other photosynthates (Burriesci et al., 2012; Davy et al., 2012; Falkowski et al., 1984; Muscatine et al., 1984). Yet this mutualism is highly temperature sensitive, as cnidarians lose their symbionts just 1–2°C above local summer mean temperatures in a process known as bleaching (Glynn, 1996; Hoegh-Guldberg et al., 2007). Bleaching is a visually striking stress response whose clear signal is an indispensable environmental monitoring tool (Hughes et al., 2017). However, recent studies have shown that heat stress can perturb coral carbon metabolism (Innis et al., 2021; Rädecker et al., 2021) and calcification (Allen-Waller and Barott, 2023; Inoue et al., 2012) even when visual bleaching does not occur. A better understanding of these more cryptic cnidarian heat stress responses is essential for predicting coral responses to climate change, and may assist in discovering additional coral resilience biomarkers (e.g., (Barshis et al., 2010)) that will help to identify stress-tolerant corals that can survive warming oceans (Putnam et al., 2017; Van Oppen and Gates, 2006).

The consequences of heat stress on the metabolism and cell physiology of marine invertebrates remain underexplored (Melzner et al., 2022), and more physiological data is necessary to predict how accelerating climate change will alter species abundance and distribution (Wagner et al., 2023). For example, thermal stress interferes with intracellular pH (pH_i_) regulation in many species. This also includes coral (Allen-Waller and Barott, 2023; Gibbin et al., 2015; Innis et al., 2021), where heat stress is threatening processes that are essential to survival and calcification, and thus reef persistence (Tresguerres et al., 2017). Understanding how heatwaves will affect cnidarian acid-base homeostasis is crucial as increasing atmospheric CO_2_ simultaneously acidifies and warms the oceans (Albright and Mason, 2013; Albright et al., 2016). However, the mechanism of thermal pH_i_ dysregulation in corals remains unclear. Disruption of symbiont CO_2_ uptake at high temperatures is one possible mechanism by which heat might lead to acidification in cells hosting endosymbionts (Gibbin et al., 2015); however, this does not explain why cells without symbionts are also more acidic after heat stress (Allen-Waller and Barott, 2023; Innis et al., 2021). ATP limitation resulting from symbiont loss might reduce available energy for acid-base homeostasis for the whole organism, as bleaching-susceptible individuals have lower pH_i_ than their bleaching-resistant neighbors during marine heatwaves (Innis et al., 2021). Yet even corals that do not bleach can suffer disrupted metabolism and pH_i_ homeostasis under heat stress (Innis et al., 2021; Rädecker et al., 2021). Symbiotic dysfunction is therefore unlikely to be the sole cause of thermal acid-base dysregulation. However, it remains unclear whether heat stress alters cnidarian pH_i_ by causing photophysiological stress in the resident Symbiodiniaceae, directly affecting host cellular processes, or by disrupting the mutualism itself.

To test the hypothesis that heat stress interferes with cnidarian acid-base regulation independent of symbiont loss, we used separate symbiotic and aposymbiotic subpopulations of a clonal line of the symbiotic sea anemone *Exaiptasia diaphana*, a robust model for cnidarian symbiosis that can live indefinitely in culture without symbionts (Sunagawa et al., 2009; Weis et al., 2008). This system allows us to disentangle the direct effects of heat stress on the host from stress resulting from bleaching, avoiding complications associated with the severe stress experienced by corals following symbiont loss at high temperatures (Brown, 1997; Davy et al., 2012; Jones, 2008; Oakley and Davy, 2018). In order to investigate host-derived changes in acid-base homeostasis in comparison with physiological disruptions resulting from bleaching, symbiotic and aposymbiotic *E. diaphana* (CC7 clone) (Sunagawa et al., 2009) from two separate cohorts were exposed to four increasing temperature treatments (target temperatures: 25°C (ambient control), 27°C, 29°C, and 31°C) for two weeks. We then measured symbiont density, protein content, intracellular pH (pH_i_), symbiont and host carbon assimilation, and metabolic rates (photosynthesis and respiration). We hypothesized that elevated temperatures would alter host pH_i_ in aposymbiotic as well as symbiotic anemones, but that higher rates of photosynthesis and photosynthate translocation to the host could mitigate pH_i_ dysregulation in symbiotic individuals. These data provide important insights into how climate change alters cellular homeostasis in the context of endosymbiosis.

## MATERIALS AND METHODS

### Anemone populations

Symbiotic and aposymbiotic *Exaiptasia diaphana* of the CC7 clonal strain were used in this study (Sunagawa et al., 2009). Two distinct CC7 populations were used: 1) a population that had been maintained in the Barott Lab at the University of Pennsylvania for several years at static temperature (25°C) and photosynthetically active radiation (PAR) of 100 µmol m^−2^ s^−1^, and 2) a population that had been maintained in the Cleves Lab at the Carnegie Institution for Science for several years (25°C; 25 µmol m^−2^ s^−1^) that was obtained three weeks prior to the experiment (Fig. 1). In both populations, aposymbiotic animals were generated from a subset of the corresponding symbiotic population and were maintained in an aposymbiotic state for several years prior to the start of the experiments. Aposymbiotic animals were kept in the dark in opaque tubs to prevent symbiont re-colonization and symbiotic anemones were kept in transparent tubs. All anemones were kept at 25°C under 100 µmol m^−2^ s^−1^ and fed freshly hatched *Artemia* weekly prior to the start of the experiment.

**Figure 1.**
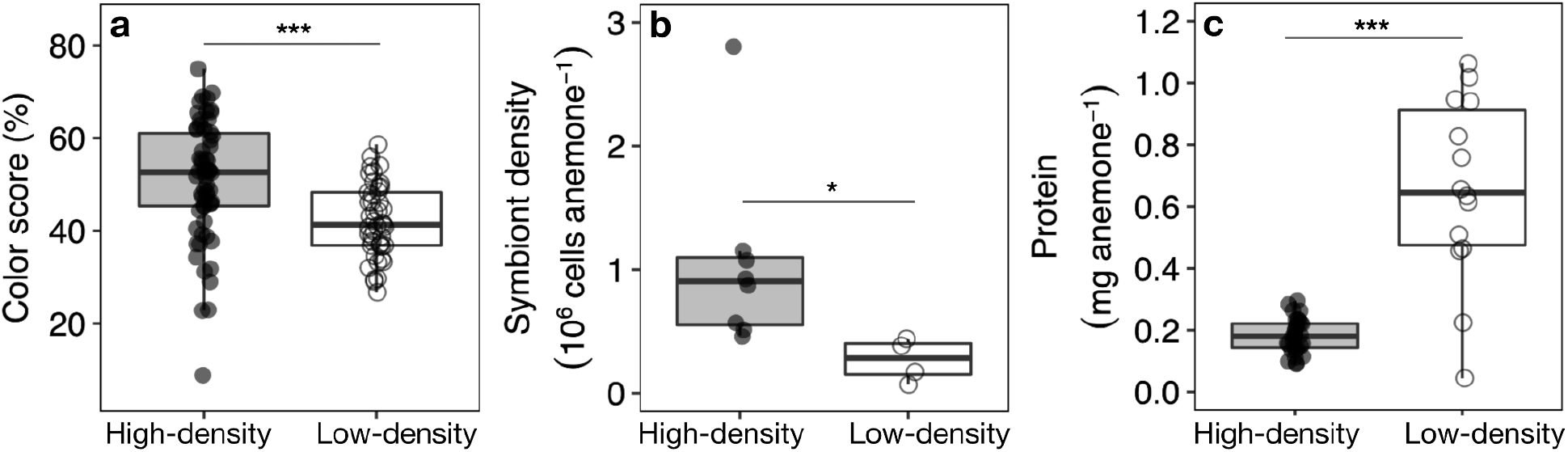
Cohorts of clonal *Exaiptasia diaphana* showed baseline differences in color, symbiont density, and protein. **a)** Pre-treatment color scores (% red color) of symbiotic anemones differed by cohort. **b)** Symbiont cells per anemone at baseline incubation temperature (25°C) differed by cohort. **c)** Protein concentration differed by cohort. Points represent individual anemones. Boxplots show medians with 25th and 75th percentiles, and whiskers show 1.5x interquartile range. Asterisks show results from linear mixed effects models with cohort as fixed effect and anemone container as a random effect (*P<0.05, ***P<0.001).

### Heat stress experiment setup and monitoring

Anemones were randomly assigned to each of the four temperature treatments for both experiments. Target temperatures (25, 27, 29, and 31°C) were chosen to encompass a range of heat above anemone culture temperature (25°C) (Fig. 2). During the experiment, the anemones were kept in 118-mL containers (N=4 containers per treatment per symbiont status for each cohort, with 3-4 anemones per container; Fig. 2a) (Ziploc Twist n Loc, SC Johnson, Racine, WI) filled with 0.22 µM-filtered artificial sea water (FSW) (salinity ∼ 34 ppt; Instant Ocean Reef Crystals, Spectrum Brands, Blacksburg, VA). Aposymbiotic and symbiotic anemones were placed in separate containers, and the exterior of all aposymbiotic containers and lids were covered with black electrical tape to maintain dark culture conditions. Containers with symbiotic anemones were covered with transparent plastic film secured with rubber bands to minimize evaporation while permitting light to pass through. All containers were set on racks in water baths (40 gal; one per temperature in each cohort) each equipped with a circulating pump (1500L/H Submersible Water Pump, Vivosun, Ontario, CA) and 1–2 50W heaters (either Aqueon Submersible Water Heater, Central Garden & Pet Company, Franklin, WI or Hygger Fish Tank Water Heater, Shenzhen Mago Trading Co., Shenzhen, China). Anemones were kept on a 12:12-hr light:dark cycle (NICREW HyperReef LED, Shenzhen NiCai Technology Co., Shenzhen, China), with photosynthetically active radiation averaging 160–175 µmol m^−2^ sec^−1^ across the duration of the experiment (Table 1). Anemones were fed *Artemia* weekly through the experiment until 1 week prior to sampling and anemone containers were cleaned every ∼3 days.

**Figure 2.**
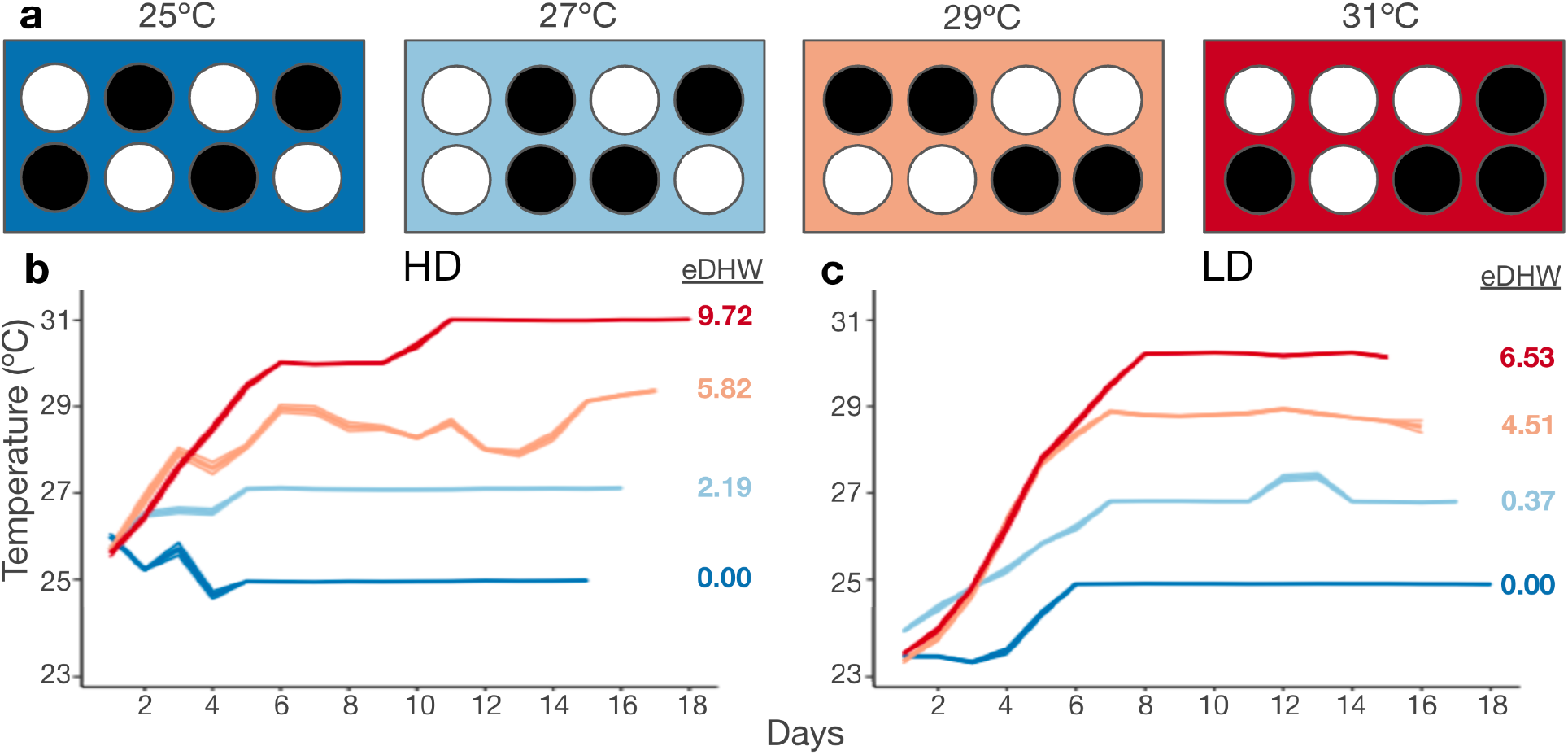
Experimental design and temperature conditions across the experimental period. **a)** *Exaiptasia diaphana* were kept in separate containers (N=4 containers per treatment for each cohort; 3-4 anemones per container) within water baths held at 25, 27, 29, or 31°C. Daily average temperatures (24 hr mean) across containers within temperature treatments of the **(b)** high-symbiont-density (HD) and low-symbiont-density (LD) **(c)** cohorts. Ribbons indicate standard error. Seawater temperatures were significantly different by treatment within each cohort (P<0.001). Annotations to the right of each temperature graph indicate total accumulated experimental degree heating weeks (eDHW; °C week^−1^) for each treatment.

**Table 1.**
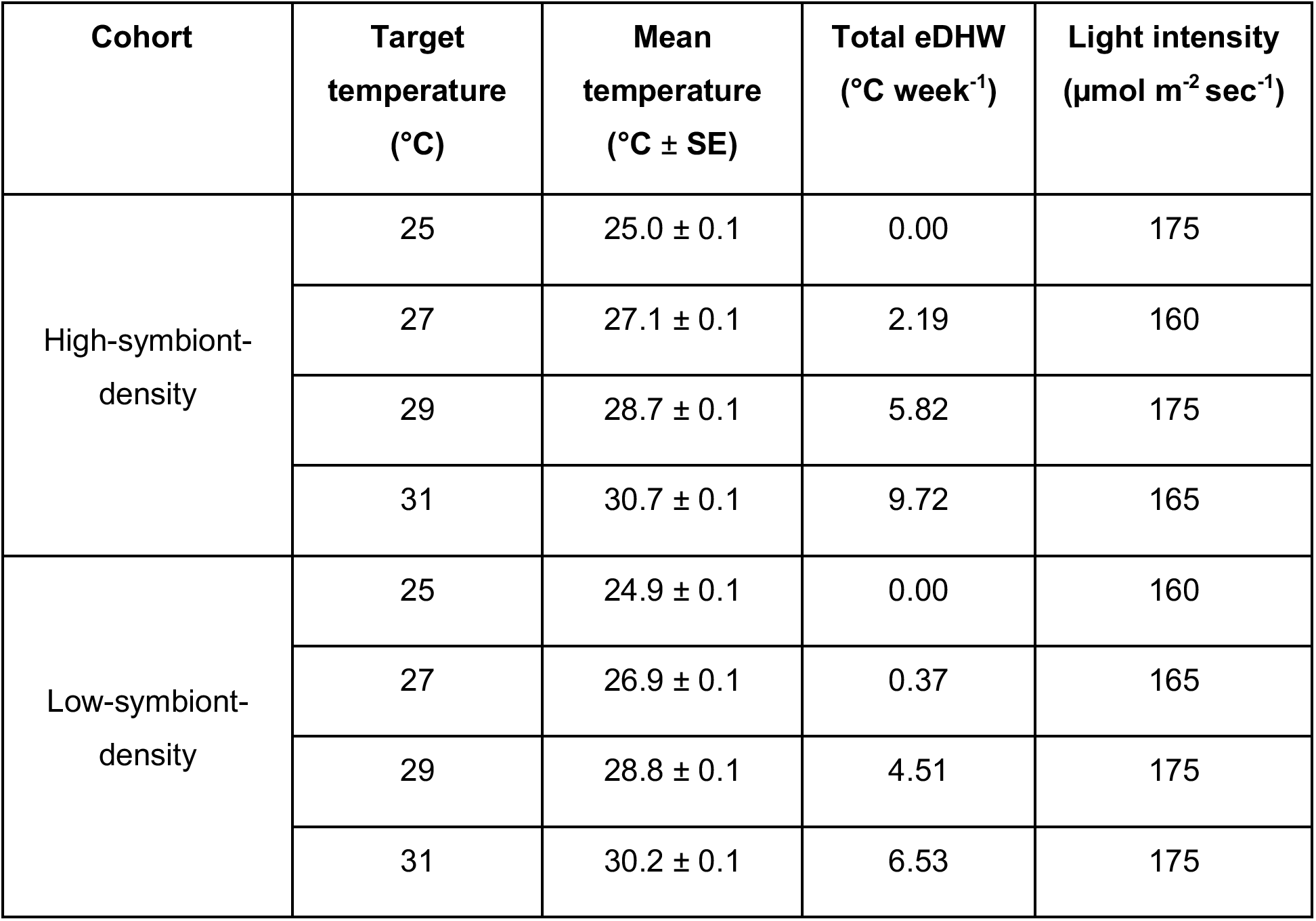
Experimental temperature treatments. SE, standard error; eDHW, experimental degree heating week.

Containers were placed inside of the tanks 2 d prior to each experiment to acclimate anemones to their new surroundings. All tanks started at 25°C on Day 0 and were heated 1°C/d until they reached their target temperatures (4-8 days; Fig. 2b-c). Seawater temperatures were recorded every ≤ 15-120s using cross-calibrated temperature loggers (accuracy: ± 0.11 to 0.29°C; HOBO UA-001-64, Onset Computer Corporation, Bourne, MA). Daily temperature measurements of the water bath were also recorded to ensure proper progression of the heat ramp (ProDSS with ODO/CT Probe, YSI, Yellow Springs, OH). In addition, temperatures inside each anemone container were measured on 6 separate days through the experiment. As containers were consistently cooler than the surrounding water bath, we calculated each treatment’s average temperature (Fig. 2b-c) based on the average difference (T_tank_ - T_container_) subtracted from the water bath temperature. Anemones were monitored daily for mortality, and photographed every other day with a color standard for color quantification. Anemone color was quantified in ImageJ (Abramoff et al., 2004) using a modified coral color quantification protocol (Allen-Waller and Barott, 2023). Briefly, each image was converted to RGB, and the red channel was further examined. An ovoid region of interest was drawn to encompass each individual’s oral disk, and intensity in that channel was divided by the intensity of the red standard in the same image to obtain percent intensity. Color scores are reported as [100% - (% red intensity)] because 100% red intensity corresponds to zero red pigment, a proxy for chlorophyll *a* concentration.

Anemones within the same treatment were sampled on the same day for each treatment temperature (one treatment per day over a four day period) for several metrics: one anemone from each container was measured for *in vivo* metabolic rates (high-density cohort only), then frozen at −80°C for symbiont and protein content analysis; one was used for *in vivo* intracellular pH measurements; and one underwent a ^13^C stable isotope pulse-chase.

### *In vivo* photosynthesis and respiration

Photosynthesis-irradiance curves were performed to determine photosynthetic and light-enhanced dark respiration (LEDR) rates. Oxygen (O_2_) evolution was measured from 0 to 270 µmol m^−2^ s ^−1^ on a subset of anemones (n = 8 of each symbiotic status) from each treatment condition following 2 weeks of exposure, totaling 32 anemones per treatment (high-density cohort only). Each anemone was individually enclosed in a 4 mL glass vial full of FSW and equipped with an optical O_2_-sensor (SV-PSt5, Presens, Regensburg, Germany). For each temperature treatment, four vials were filled with FSW and measured alongside anemones as blanks. Vials were then placed in a light-masked sensor dish reader (SDR SensorDish Reader with SDR-MSV24, Presens) that optically measured O_2_ concentration in 24 vials simultaneously every 15 s while being mixed continuously at 200 rpm on a plate shaker (Talboys Standard Orbital Shaker, Troemner, West Deptford Township, NJ) throughout the incubation. For each treatment group, O_2_ evolution was measured for ≥ 10 min at each of six increasing light levels (24, 60, 113, 165, 217, and 270 µmol m^−2^ s^−1^), then in total darkness for ≥ 10 min. Measurements were taken inside of an incubator (B.O.D. Low Temperature Refrigerated Incubator, VWR, Radnor, PA) set to each group’s experimental temperature, and constant temperatures during incubation were verified using a temperature logger (accuracy: ± 0.11 to 0.29 °C; HOBO UA-001-64, Onset Computer Corporation) placed in a water bath inside the incubator. Temperature- and light-specific blank values were determined by averaging O_2_ evolution across the FSW-only vials within each light level, and this value was then subtracted from each anemone’s measured O_2_ evolution rate to account for microbial O_2_ evolution. A total of 12 individuals (1 symbiotic and 11 apo) had oxygen evolution rates within ± 1 standard error of blank values in the dark, and were excluded from downstream analyses as they were deemed below the limit of detection, likely as a result of their small body size (55–140 µg protein anemone^−1^). In total, N=15 symbiotic and N=5 aposymbiotic anemones were used for subsequent metabolic analyses.

### Symbiont density

Anemones were thawed and homogenized in 500 µL deionized H_2_O at 25,000 rpm for 10 s using a tissue homogenizer (Fisherbrand 850 Homogenizer, Thermo Fisher Scientific, Waltham MA, USA) followed by needle-shearing until homogenous using a 22-gauge needle. Tissue homogenates were spun at 7,000 x g for 5 min to separate host (supernatant) and symbiont (pellet) fractions. Symbiodiniaceae were resuspended in 1% sodium dodecyl sulfate in 1X phosphate-buffered saline and cell concentrations were determined in triplicate using a flow-cytometer (Guava easyCyte 5HT, Luminex, Austin TX, USA) as described (Innis et al., 2021; Krediet et al., 2015). Protein concentration in the host fraction was measured on a spectrophotometer (BioTek PowerWave XS2, Agilent, Santa Clara CA, USA) using Coomassie Plus Bradford assay reagent (Pierce, Thermo Fisher Scientific).

### Intracellular pH

Cells were isolated from one randomly selected anemone from each anemone container (n = 4 of each symbiotic state per treatment) by holding the anemone with forceps and brushing it against a small soft-bristled toothbrush submerged in 15 mL FSW for 2–4 min to break down tissue until enough isolated cells were obtained. The resulting cell suspension was passed through a 40 µM cell strainer (Fisherbrand). Cells were then spun at 350 x g for 4 min and resuspended in 1 mL FSW with 10 µM pH-sensitive cell dye SNARF-1 acetoxymethyl ester acetate (Invitrogen, Thermo Fisher Scientific) and 0.1% dimethylsulfoxide (Invitrogen) as described (Venn et al., 2009) for 20 min at 25°C in the dark. Live cells were then pelleted to remove any dye that did not enter the cells, resuspended in 1 mL FSW, and imaged at 25°C in a glass-bottomed dish using an inverted confocal microscope (Leica SP8 DMi8, Leica Camera, Wetzlar, Germany) at 63x magnification (HC PL Apochromat C52 Oil objective, numerical aperture = 1.4). All samples were excited at 514 nm (10% power argon laser, 1% emission, 458/514/561 beam splitter) and SNARF-1 fluorescent emission was simultaneously acquired in two channels (585 ± 15 and 640 ± 15 nm) using HyD detectors (gain = 100, pinhole = 1.00 airy unit). 512 x 512 pixel images were acquired at 8 bits/pixel using a scan speed of 400 Hz. Between 9–22 gastrodermal cells containing symbionts (symbiocytes) and 7–38 cells without symbionts (non-symbiocytes) were measured per symbiotic anemone; there were no symbiocytes present in aposymbiotic anemones so only non-symbiocytes are reported. SNARF-1 fluorescence ratios in anemone cytoplasm were quantified using ImageJ and normalized to background fluorescence as described (Innis et al., 2021), and converted to pH values using a calibration curve generated on the same microscope within 2 weeks of the experiment as described (Venn et al., 2009) (Fig. S1).

### Stable isotope tracer experiment

One anemone per symbiotic anemone container (n = 4 per treatment) was randomly selected to measure fixed carbon assimilation in the symbiont and translocation to the host using a stable isotope (NaH^13^CO_3_) pulse-chase experiment. In addition, three wild-type aposymbiotic anemones were included during the 25°C incubation as controls, expected to show no ^13^C assimilation due to a lack of symbionts. At 11:00 each sampling day, single anemones were placed in separate 50-mL conical tubes (Corning Falcon, Fisher Scientific) filled with FSW enriched with 31.22 µM NaH^13^CO_3_ (Cambridge Isotope Laboratories, Andover, MA). Anemones were then replaced in their experimental temperature and light conditions inside their sealed tubes, upside down in a tube rack submerged in their water bath, for a 7-hr ‘pulse’ period so that symbionts could incorporate ^13^C via photosynthesis. After 7 h (18:00), the pulse was removed, anemones were rinsed with unamended FSW, and tubes were refilled with unamended FSW for the 12-hr ‘chase’ period. At 06:00 the following morning, the FSW was removed and anemones were frozen at −80°C and maintained at −80°C until processing. Samples were then thawed, homogenized, and separated into host and symbiont fractions as described above. Each fraction was then transferred directly into a pre-weighed opened tin capsule (EA Consumables, Marlton, NJ) and dried to stable weight for 24 h at 50°C. Capsules were then closed, weighed, and shipped to the University of California, Santa Cruz Stable Isotope Facility (Santa Cruz, CA) for ^13^C enrichment analysis by elemental analysis (NC2500 Elemental Analyzer, Carlo Erba Reagents GmbH, Emmendingen, Germany) coupled with isotope ratio mass spectrometry (Delta Plus XP iRMS, Thermo Scientific; coupling via: Conflo III, Thermo Scientific).

### Statistical analysis

All data were analyzed in RStudio version 2022.07.2 (https://www.rstudio.com/) and plots were generated using the package *ggplot2* (Wickham, 2016). To find the best fit model for each response variable, relevant linear, linear mixed effects, and generalized additive models were developed using the *lme4* and *mgcv* packages (Bates; Wood, 2011). Corrected Akaike Information Criterion (AICc) were calculated for each model using the *MuMIn* package (Barton 2015) and the model with the lowest AICc was chosen. Q-Q and residual plots were checked to ensure each model met normality and homogeneity of variance assumptions. All models are summarized in Table S1.

#### Temperature treatments

To compare temperatures of treatments within each cohort, one linear model analyzing daily average (24 hr mean) container temperature with treatment as a fixed effect (four levels: 25°C, 27°C, 29°C, and 31°C) was constructed for each cohort (HD and LD). To compare treatments between the two cohorts’ experiments, a linear model analyzing daily average container temperature with fixed effects of temperature (four levels: 25°C, 27°C, 29°C, and 31°C) and cohort (two levels: HD and LD) was constructed. Significant interactions were further explored with Tukey’s HSD adjusted post-hoc pairwise comparison tests using the *emmeans* package (Lenth and Lenth, 2017). Experimental degree heating weeks (eDHW) were calculated for each treatment by summing the absolute value of the difference between the daily average and 25°C, for all days over the course of the experiment where the daily average exceeded 26°C (mean monthly maximum + 1°C (Leggat et al., 2022)). Mean monthly maximum was designated as 25°C since anemones were all maintained at 25°C before the experiment.

#### Cohort physiological comparison

The two experimental cohorts’ symbiont densities were assessed by comparing 25°C control anemones from both cohorts. Temperature treatment was not found to have an effect on protein, so cohort protein densities were compared using protein content measurements of all physiology samples from each cohort. To verify that the pulsed ^13^C isotope was enriched in symbiotic anemones of both cohorts, carbon assimilation was compared between symbiotic host tissue, symbionts, and aposymbiotic host control tissue using a linear model with effects of cohort (two levels: HD vs. LD), symbiont status (two levels: symbiotic and aposymbiotic), and tissue fraction (two levels: host and symbiont).

#### Physiological response to temperature

Since cohorts differed in initial symbiont density and biomass, they were tested separately for the effect of temperature treatment (four levels: 25°C, 27°C, 29°C, and 31°C) and symbiotic status (two levels: symbiotic and aposymbiotic) on anemone physiology (host protein, symbiont density, symbiont carbon assimilation, host carbon assimilation, symbiocyte intracellular pH, non-symbiocyte intracellular pH). For each variable, the best model was selected according to lowest AICc, after which any significant effects were further explored with Tukey’s HSD adjusted post-hoc pairwise comparison tests using the *emmeans* package (Lenth and Lenth, 2017). Notably, refined models of non-symbiocyte intracellular pH that did not consider symbiotic status all had lower AICc than full models that accounted for symbiotic status, so aposymbiotic and symbiotic animals were combined for subsequent analysis of pH_i_ in that cell type.

#### Respirometry

To test effects of irradiance on O_2_ evolution, generalized additive models (GAMs) with light level as a fixed effect were fit separately for each HD cohort symbiotic and aposymbiotic anemone from each temperature using a modified respirometry analysis procedure (Becker and Silbiger, 2020). The maximum oxygen evolution rate from each anemone’s predicted photosynthesis-irradiance GAM fit was calculated and defined as that anemone’s maximum photosynthetic rate (P_max_). The optimal temperature for both P_max_ and light-enhanced dark respiration (LEDR) was then found by fitting secondary GAM fits for P_max_ and LEDR over all experimental temperatures and calculating the inflection point using the first derivative of the GAM spline. Best-fit GAMs for P_max_ and LEDR response to temperature were selected as described above.

#### Principal components analysis

For symbiotic animals only, differences in physiology were analyzed separately for each cohort using permutational multivariate analysis of variance (PERMANOVA) using the adonis function of the *vegan* package (Oksanen et al., 2013) and principal components analysis (PCA) using the R *stats* package prcomp function with temperature as a fixed effect. To assess differences between anemones with different symbiont densities, a subset of all physiological response variables that were measured in all anemones (protein content, symbiont density, and non-symbiocyte intracellular pH) was used to compare all individuals using a separate PERMANOVA and PCA with cohort (two levels: HD vs. LD) and symbiotic status (two levels: symbiotic vs. aposymbiotic) as fixed effects.

## RESULTS

### Anemone cohort traits

Symbiotic anemones from the two experimental cohorts differed significantly in symbiont density (F=3.96, P=0.047) and color (F=13.86, P<0.001) at control temperatures (Fig. 1). Specifically, symbiotic anemones from one cohort were darker in color than the other (Fig. 1a) and hosted more symbionts per anemone (Fig. 1b); thus, they were termed high-symbiont-density or HD anemones. Symbiotic anemones from the other cohort were paler (Fig. 1a) and contained fewer symbionts (termed low-symbiont-density or LD) (Fig. 1b). Symbiotic LD anemones also contained more protein per anemone than symbiotic HD anemones (F=75.94, P<0.001; Fig. 1c), corroborating visual observations of larger body size in the LD cohort. Further, for both HD (F=50.0, P<0.001) and LD (F=57.7, P<0.001) cohorts, aposymbiotic anemones were significantly smaller and had fewer symbionts than their symbiotic counterparts (Fig. S2).

### Experimental conditions

Seawater temperatures in containers from all four treatments differed significantly from one another within both the HD (F=528.68, P<0.001; Tukey’s HSD: P<0.001 for all pairwise comparisons) and LD (F=2682.3, P<0.001; Tukey’s HSD: P<0.001 for all pairwise comparisons) cohort experiments. Between the two cohorts, the lowest three temperature treatments (25, 27, 29°C) did not differ (Tukey’s HSD: P>0.15), whereas the highest temperature was an average of 0.5°C warmer in the HD cohort than the LD cohort (Tukey’s HSD: P=0.017), reaching a maximum daily average of 30.7°C and 30.1°C, respectively (Table 1; Fig. 2b-c). Total experimental degree heating weeks (eDHW) for the highest temperature treatments were 9.72°C wk^−1^ for the HD and 6.53°C wk^−1^ for the LD cohort (Table 1; Figure 2b-c). Intermediate temperatures experienced eDHW ranging from 0.37–5.82°C wk^−1^, and 25°C controls in both cohorts experienced 0°C wk^−1^(Table 1; Figure 2b-c).

### Temperature effects on symbiont density and photosynthate assimilation

Temperature differentially impacted the two anemone cohorts. As temperatures increased, HD anemones showed a significant decline in symbiont density (F=4.86, P=0.005; Fig. 3a; Fig. S2) and a marginally insignificant trend of less symbiont photosynthate assimilation (F=3.00, P=0.072; Fig. 3c), which resulted in anemones experiencing the highest temperatures to assimilate less photosynthate than controls (T=3.04, P=0.044; Fig. 3e). By contrast, heat did not cause significant reductions in symbiont density in LD anemones (F=0.06, P=0.98; Fig. 3b). Symbiont photosynthate assimilation in LD anemones was higher than in HD anemones (T=-4.39, P<0.001; Fig. S3) and although LD assimilation was significantly affected by temperature (F=9.49, P=0.003; Fig. 3d), higher temperatures did not consistently decrease host photosynthate assimilation (F=1.75, p=0.22; Fig. 3f). In fact, symbiont loss was only correlated with lower host photosynthate assimilation in HD anemones, and the correlation was weak (F=5.54, P=0.034, R^2^=0.23; Fig. S4a). In LD anemones, anemones with fewer symbionts did not assimilate less photosynthate (F=0.12, P=0.73, R^2^=-0.07; Fig. S4b). Across both cohorts, aposymbiotic anemones assimilated significantly less ^13^C than symbiotic hosts (F=-2.11, P=0.040), which in turn contained significantly less ^13^C than their symbiont populations (F=-10.56, P<0.001), confirming that the ^13^C pulse successfully enriched photosynthates in symbiont and host fractions (Fig. S3).

**Figure 3.**
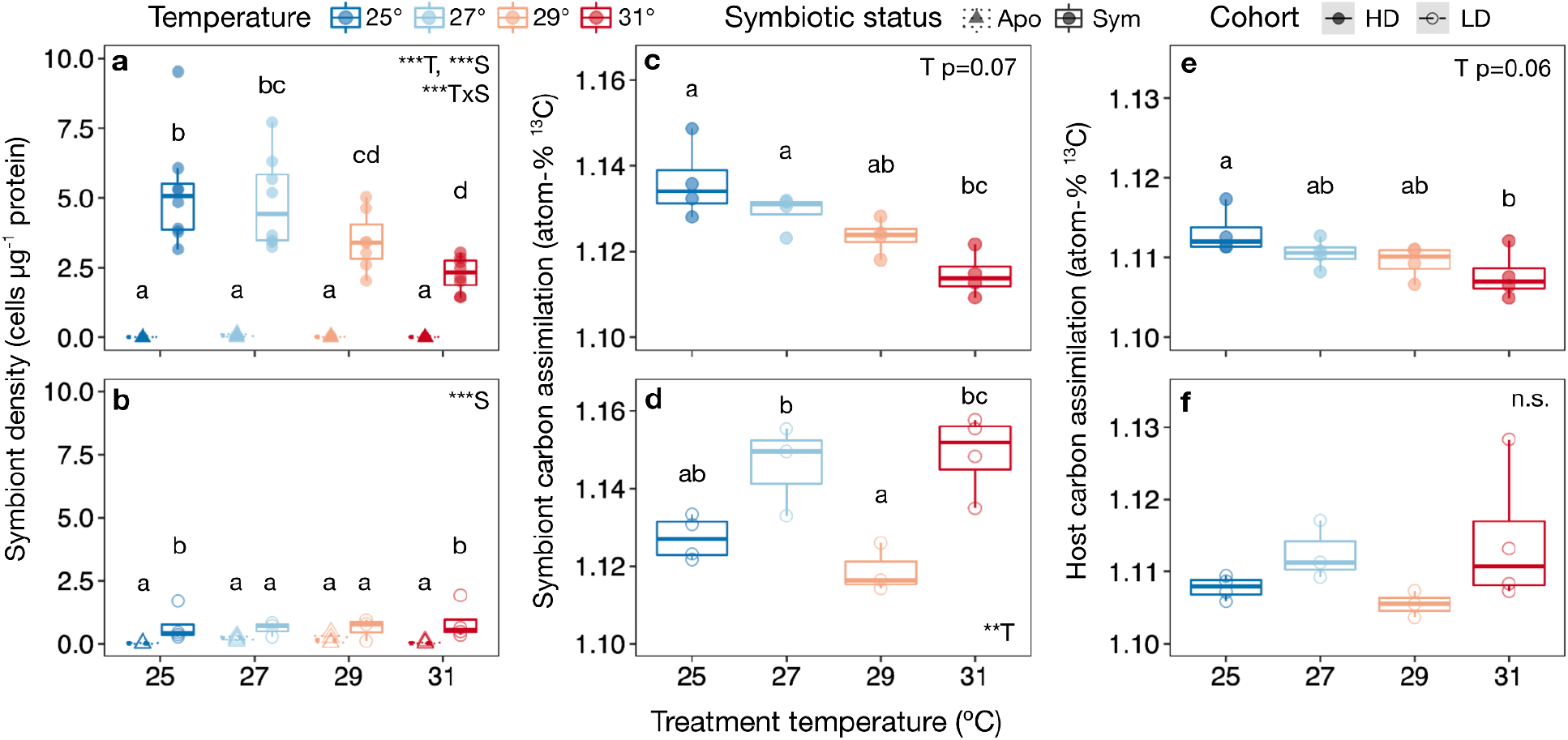
Temperature altered dinoflagellate density and symbiotic function in high-symbiont-density but not low-symbiont-density *Exaiptasia diaphana*. **a-b)** Symbiont density response to temperature density in symbiotic (sym, circles) and aposymbiotic (apo, triangles) individuals differed between high-symbiont-density (HD) **(a)** and low-symbiont-density (LD) **(b)** anemone cohorts. **c-f)** Temperature response of symbiont **(c-d)** and host **(e-f)** photosynthate assimilation (atom-% ^13^C) differed between **(c, e)** high-symbiont-density and **(d, f)** low-symbiont-density anemones. Points represent individual anemones. Boxplots show medians with 25th and 75th percentiles, and whiskers show 1.5x interquartile range. Insets show results of linear models with effects of temperature (T), symbiotic status (S), and their interactions (*P<0.05, **P<0.01, ***P<0.001). Small letters denote significant pairwise groupings (P<0.05) (Tukey’s HSD).

There was no effect of temperature on protein content in symbiotic or aposymbiotic anemones from either the HD (F=0.433, P=0.73) or LD cohort (F=0.312, P=0.816) (Fig. S3a-b). Symbiont density model results were consistent whether symbiont density was calculated as cells per animal or per anemone protein (Fig. S3c-d), confirming that the observed changes in symbiont density per protein were not a result of changes in biomass. Two symbiotic anemone containers (one container each from the 27°C and 29°C treatment; six anemones total) from the LD cohort experienced total mortality on day 14 and were not analyzed.

### Anemone metabolic performance

Photosynthesis:irradiance curves, measured only on the HD cohort of anemones, showed significant metabolic changes across light levels in symbiotic anemones (edf: 1.98, F=70.77, P<0.001) and aposymbiotic anemones (edf: 1.94, F=11.83, P<0.001), with the fastest photosynthesis and light-enhanced dark respiration (LEDR) rates observed in the 27°C-incubated symbiotic group (Fig. S5a-b). Generalized additive models (GAM) fit to Pmax and LEDR values for symbiotic individuals across the temperature treatments confirmed that temperature significantly affected both photosynthesis (edf: 2.79, F=4.22, P=0.021) and LEDR (edf: 2.33, F=4.46, P=0.030) in symbiotic anemones (Fig. 4). The predicted optimal temperature for Pmax was 26.39 ± 1.39°C (Fig. 4a), and the predicted temperature for maximum LEDR was 27.23 ± 0.57°C (Fig. 4b). In aposymbiotic anemones, temperature significantly influenced LEDR (F=14.22, P=0.001) but not Pmax (F=3.19, P=0.090) (Fig. S5c-d).

**Figure 4.**
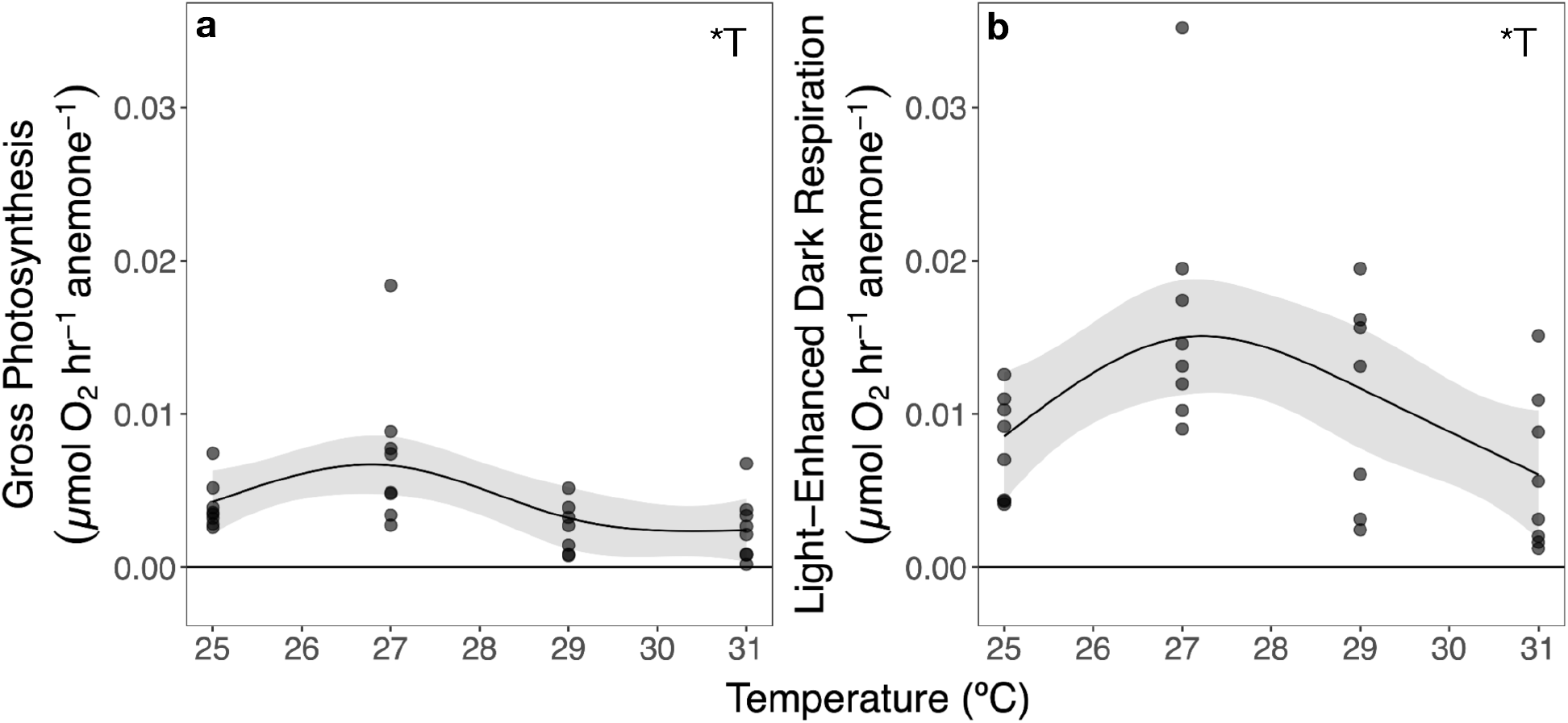
*Exaiptasia diaphana* metabolic rates vary with temperature. Maximum gross photosynthesis **(a)** and respiration **(b)** of high-symbiont-density symbiotic anemones after incubation at treatment temperatures. Points represent model-estimated metabolic rates for individual anemones. Lines indicate metabolic rates predicted by generalized additive models for temperature (T) (gray ribbon = 95% confidence interval), with insets indicating significant effects (*P<0.05).

### Effect of elevated temperatures on anemone intracellular pH

Temperature had a significant effect on intracellular pH (pHi) for cells from both aposymbiotic and symbiotic anemones (Fig. 5). Symbiont-hosting cells (symbiocytes) tended to become more acidic with increasing temperature. This effect was stronger in the LD (F=4.25, P=0.035, Fig. 5c) than in the HD cohort (F=3.00, P=0.073, Fig. 5a). Cells without symbionts (non-symbiocytes) showed a significant but non-monotonic temperature response (HD: F=10.97, P<0.001; LD: F=10.22, P<0.001, Fig. 5d), with 27°C-treated anemones having the most acidic cells in both cohorts (Fig. 5b,d). Within each cohort, there was no effect of anemone symbiotic state on pHi: non-symbiocytes from aposymbiotic anemones had the same pHi as non-symbiocytes from symbiotic anemones at each temperature, and showed the same thermal acid-base disruption as non-symbiocytes from symbiotic anemones (Fig 5b,d). Interestingly, non-symbiocyte pHi differed between cohorts, and was significantly higher in the HD cohort than the LD cohort across temperatures (F=22.08, P<0.001; Fig. 5f). Moreover, host photosynthate assimilation only correlated with pHi of symbiocytes from HD anemones (F=7.98, P=0.013, R^2^=0.32; Fig. 5e); there was no association between host photosynthate and pHi of HD non-symbiocytes (F=0.056, P=0.82, R^2^=-0.07; Fig. 5f) or of either cell type in LD anemones (non-symbiocytes: F=0.76, P=0.40, R^2^=-0.02; symbiocytes: F=1.62, P=0.22, R^2^=0.05; Fig. 5e-f).

**Figure 5.**
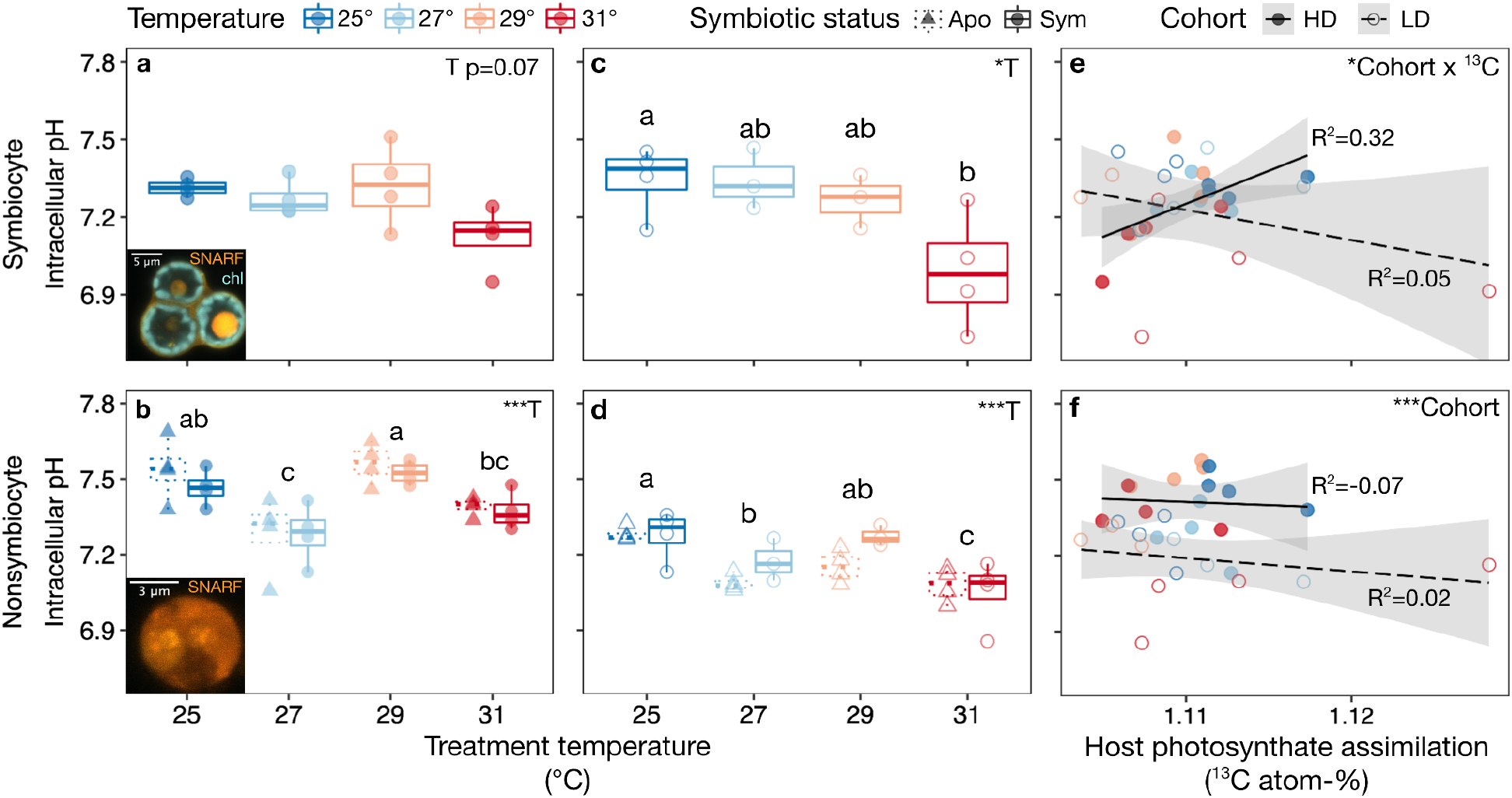
Heat altered *Exaiptasia diaphana* intracellular pH (pHi_i_) regardless of symbiont presence, density, or mutualistic function. **a-b)** pH_i_ of high-symbiont-density (HD) *E. diaphana* symbiocytes **(a)** and non-symbiocytes **(b)** varied with temperature. Insets, symbiocyte **(a)** and non-symbiocyte **(b)** showing fluorescence of pH-sensitive dye (SNARF-1) and symbiont chlorophyll (chl)**. c-d)** pH_i_ of low-symbiont-density (LD) *E. diaphana* symbiocytes **(c)** and non-symbiocytes **(d)** decreased with temperature. Inset capital letters show results of linear models with effects of temperature (T); best-fit models found no effect of anemone symbiotic status (symbiotic = Sym, circles; aposymbiotic = Apo, triangles). Small letters denote significant pairwise groupings (P<0.05) (Tukey’s HSD). Points represent individual anemones. Boxplots show medians with 25th and 75th percentiles, and whiskers show 1.5x interquartile range. **e)** In HD anemones, host photosynthate assimilation was correlated with symbiocyte pH_i_. In LD anemones, there was no correlation. **f)** Host photosynthate assimilation did not affect non-symbiocyte pH_i_ in HD or LD anemones. Points show individual anemones. Regression lines show linear models with effects of host photosynthate assimilation (^13^C), cohort, and their interaction; gray band represents 95% confidence interval; and insets show results of linear models (*P<0.05, ***P<0.001).

### Physiological separation by temperature and symbiotic status

Principal components analysis revealed symbiotic anemone physiology differed by temperature in both cohorts (HD: F=4.17, P=0.001; LD: F=2.12, P=0.026; Fig. S6). This consistent effect was largely driven by differences in intracellular pH and photosynthate assimilation (Fig. S6). However, separation of temperature groups differed between cohorts, largely because HD anemones in the hottest treatment assimilated less photosynthate than 25°C controls (Fig. S6a, Fig. 3c-d), while LD anemones under the same conditions assimilated more photosynthate than controls (Fig. S6b; Fig. 3e-f). Anemones also clustered separately by both cohort and symbiotic status: consistent with their difference in overall temperature response, anemones from the two cohorts and symbiotic statuses all separated by host protein content and non-symbiocyte pHi, as well as by symbiont density (Fig. 6).

**Figure 6.**
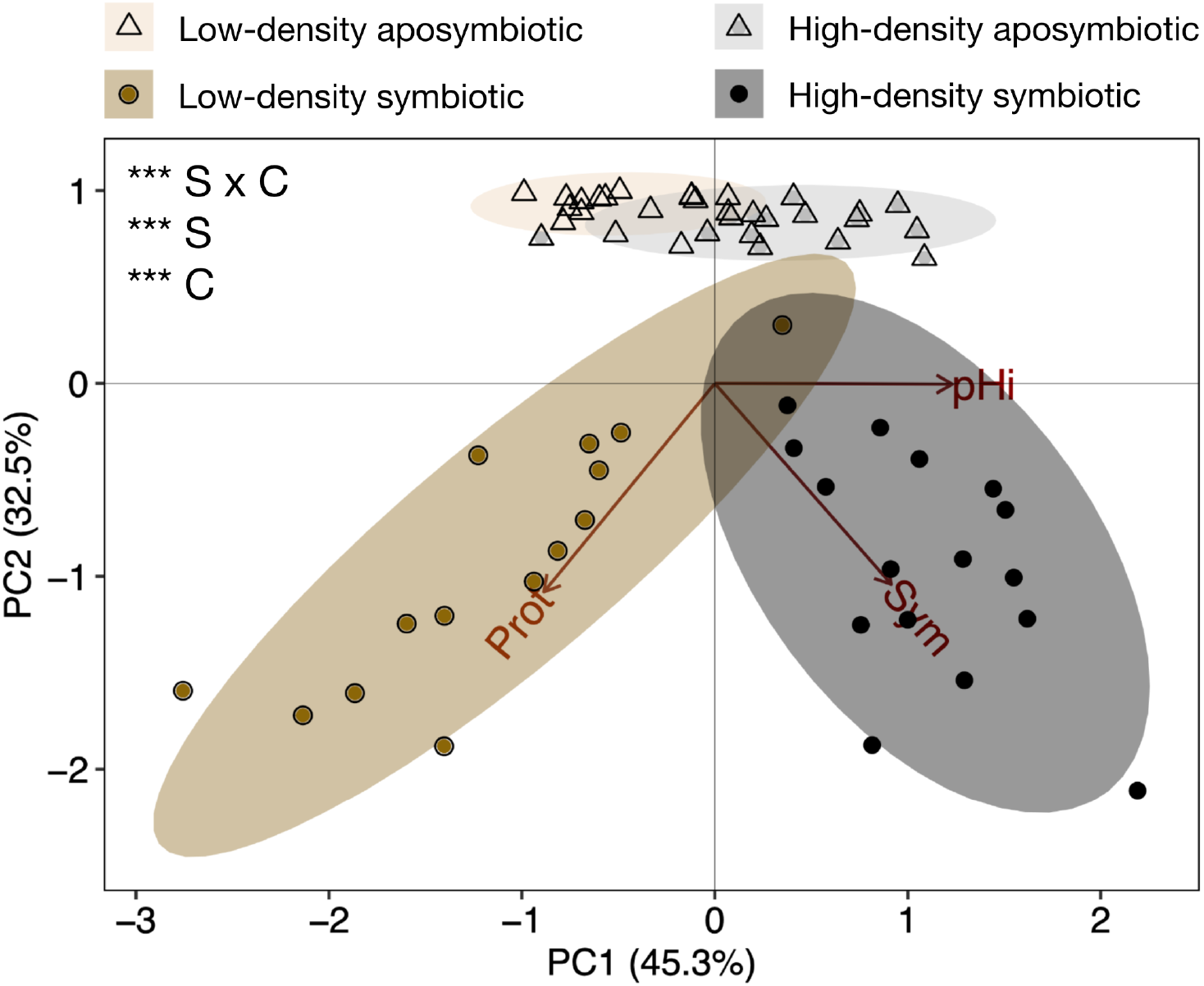
*Exaiptasia diaphana* physiology differed by degree of symbiont colonization. Simplified principal components analysis for variables shared across all experimental replicates (Prot = protein, Sym = symbiont density, and pHi = nonsymbiocyte intracellular pH) grouped by cohort and symbiotic status. Insets show significant PERMANOVA effects of cohort (C), symbiotic status (S), and their interaction (S x C) (***P<0.001). Ellipses show 95% confidence intervals for symbiotic status x cohort groupings.

## DISCUSSION

### Elevated temperature causes intracellular acidosis regardless of organismal symbiotic status

Elevated temperatures consistently led to lower intracellular pH (pH_i_) in aposymbiotic and symbiotic *Exaiptasia diaphana* relative to animals at ambient temperatures. Furthermore, symbiotic anemone cohorts exhibited similar patterns of cellular acidification at elevated temperatures, regardless of differences in initial physiologies or bleaching responses. The consistency of these responses across different anemone populations indicates that neither endosymbiont presence nor their loss is necessary for acid-base dysregulation under heat stress. While previous research in corals has suggested that heat stress alters pH regulation by interfering with symbiont photosynthesis and/or photosynthate translocation to the host (e.g. (Allen-Waller and Barott, 2023; Cameron et al., 2022; Gibbin et al., 2015; Innis et al., 2021)), the anemone responses recorded here indicate that heat stress can also impair cnidarian acid-base regulation independently of bleaching and the functional breakdown of the cnidarian-dinoflagellate symbiosis. These conclusions are not mutually exclusive, as heat stress likely disrupts both symbiont influence on and host control over host acid-base homeostasis. Multiple simultaneous host- and symbiont-derived pH regulatory mechanisms could help explain why heatwaves can impair coral acid-base homeostasis even in corals that do not lose their symbionts (Allen-Waller and Barott, 2023; Innis et al., 2021). Interestingly, bleaching resistance reduces the effect of heat on pH_i_ in corals (Innis et al., 2021), supporting contributions from both symbiont and host to acid-base dysregulation under heat stress.

Many aquatic animals experience intracellular acidosis at high temperatures (e.g., (Malan et al., 1976; Pörtner et al., 1999; Reeves and Malan, 1976)). Some of this is likely driven by a passive physicochemical effect of temperature on intracellular buffer molecules, whereby bicarbonate, phosphate, and proteins’ pK values change with temperature (Van Dijk et al., 1997). Heat may also lead to cellular acidification by decreasing lysosomal membrane stability, causing H^+^ to leak from acidic lysosomes (e.g., (Dimitriadis et al., 2012; Strand et al., 2017)). These mechanisms may have contributed to lowered cellular pH_i_ observed across *E. diaphana* cell types and symbiont densities at 31°C in the present study, and warrant further characterization in cnidarians. In addition, marine invertebrates experience increased metabolic demands at high temperatures (e.g., (Brockington et al., 2001)), including symbiotic cnidarians (Rädecker et al., 2021), and we hypothesize that this directly limits cellular capacity to regulate acid-base homeostasis. In support of this hypothesis, we observed significantly lower respiration and photosynthetic rates in symbiotic anemones at our highest treatment temperatures, suggesting metabolic impairment that could substantially reduce available energy. Thermal energy limitations can reduce pH_i_ in marine invertebrates by reducing available ATP (Pörtner et al., 1999); in cnidarians, lower ATP levels would constrain the capacity of membrane-bound ATPases and related transporters (e.g. vacuolar H^+^-ATPase, Na^+^-K^+^-ATPase, Na^+^-H^+^ exchangers) that are essential for active pH regulation in different cellular compartments (e.g., (Barott et al., 2015a; Barott et al., 2015b; Barott et al., 2022; Tresguerres et al., 2017)). Alternatively, cnidarians might respond to energy limitations by reducing expression of acid-base regulatory proteins (e.g., ion channels and transporters), as has been shown in reef-building corals (Bernardet et al., 2019; Kenkel et al., 2013). Further research is necessary to determine the relative contribution of these different possible mechanisms regulating cnidarian acid-base homeostasis.

### Bleaching response varied with symbiont density and body size

Anemone bleaching response varied with initial symbiont density, as anemones with high initial symbiont densities (HD; mean=5,233 cells µg protein^−1^) lost symbionts under heat stress, while anemones initially hosting lower symbiont densities (LD; mean=701 cells µg protein^−1^) did not. This result is consistent with greater bleaching sensitivity in cnidarians with denser symbiont loads (Cunning and Baker, 2013). The two anemone cohorts also differed in their initial biomass, which also may have influenced their bleaching responses, as the smaller-bodied HD cohort experienced significant bleaching while the larger LD cohort did not. Greater biomass may have provided the LD cohort with more nutrient resources to cope with symbiont stress at high temperatures: fewer symbionts in larger anemones would lead to a higher host biomass:symbiont ratio, which could have prevented overloading of reactive oxygen species from symbiont dysfunction during heat stress (Cunning and Baker, 2013; Nesa and Hidaka, 2009; Wooldridge, 2009). Differences in biomass may also suggest divergent trophic strategies between the two cohorts: the LD cohort had greater biomass despite hosting fewer symbionts, implying these anemones relied more on heterotrophy than the HD cohort (Hoogenboom et al., 2015). In reef-building corals, higher heterotrophic feeding can compensate for symbiont loss (e.g. (Grottoli et al., 2006)) and can increase symbiont photosynthate assimilation (Krueger et al., 2018) and translocation from symbionts during heat stress (Tremblay et al., 2016). Higher-biomass anemones may have also had more energy reserves (e.g., carbon-rich lipids) to catabolize in the absence of autotrophic resources. Conversely, reliance on autotrophy in HD anemones, evidenced by a weak but significant correlation between symbiont density and host photosynthate assimilation, suggests bleaching more negatively affected these anemones than those that were more trophically plastic.

Heterotrophy substantially affects organismal physiology in *E. diaphana*: higher host heterotrophy can increase symbiont photosynthesis (Cook et al., 1992), though greater symbiont productivity does not guarantee better host growth (Muller-Parker, 1985). This may help explain why HD anemones did not have larger body sizes despite hosting more symbionts. However, symbiont density did significantly modulate holobiont responses to heat stress. Symbiotic anemones from the lowest and highest temperatures separated in principal components analysis consistently across cohorts, driven largely by changes in carbon assimilation as well as pH_i_. Differences in carbon assimilation across temperatures were expected, given that heat stress decreases Symbiodiniaceae carbon assimilation (Ros et al., 2020) and translocation to cnidarian hosts (Allen-Waller and Barott, 2023; Baker et al., 2018; Tremblay et al., 2016). Yet heat stress only depressed host photosynthate assimilation in anemones with denser initial symbiont populations, and the HD cohort assimilated less photosynthate overall. The mechanisms driving this pattern are unknown, but could include limited light, CO_2_, or nutrients at high symbiont densities *in hospite* (Cunning et al., 2015a; Cunning et al., 2017; Hoogenboom et al., 2010; Wooldridge, 2009). However, HD anemones’ higher photosynthate assimilation than LD anemones under control conditions (25°C), along with their greater thermal sensitivity of carbon assimilation, are consistent with the hypothesis that HD anemones relied more on autotrophic energy at permissive temperatures, rendering them more likely to bleach when temperatures increased. These cohort-dependent physiological differences within a clonal line of *E. diaphana* highlight the importance of repeating studies in different populations and accounting for cohort variation in this cnidarian model.

### Endosymbiosis modulates cellular response to heat stress

Symbiont presence in anemone cells modulated how host acid-base homeostasis was affected by heat stress: symbiocytes acidified progressively with increasing temperature, whereas non-symbiocytes showed a dip in pH_i_ at 27°C relative to both cooler and warmer temperatures, and then decreased again at the highest temperature. The decline in pH_i_ of non-symbiocytes at 27°C corresponded with anemones’ peak metabolic rates, suggesting that metabolism may be important to the cell types’ varying response to temperature. Specifically, excess CO_2_ from high respiration rates at this temperature could acidify non-symbiocytes, but in symbiocytes, rapidly photosynthesizing symbionts at the same temperature may have mitigated this respiratory acidosis by drawing down dissolved inorganic carbon from the cytosol (Gibbin and Davy, 2014; Gibbin et al., 2014; Laurent et al., 2013; Putnam et al., 2017). While all anemones in this experiment were dark-acclimated for at least 40 min prior to imaging to reduce the effects of photosynthesis, it is possible that anemones with more photosynthetically active symbionts had accumulated less CO_2_ prior to dark acclimation, leading to a higher cell pH_i_ that persisted even after dark-acclimation. In addition, symbionts may have still been able to sequester CO_2_ in the dark, since symbionts isolated from *E. diaphana* express phosphoenolpyruvate carboxylase (Tytler and Trench, 1986).

Cell types may also have varied in their acidosis response due to differences in cytosolic buffering capacities between cell lineages (Madshus, 1988). Since non-symbiocytes are defined simply as cells not containing Symbiodiniaceae, they could be of any host cell lineage, while symbiocytes are a single cell type originating from the gastrodermis (Glider et al., 1980). Non-symbiocytes and symbiocytes therefore likely have pH_i_ regulatory differences simply because cnidarian cell types differ greatly in cell contents and membrane-bound transporters (Levy et al., 2021). For example, dissolved inorganic carbon transport is a major factor affecting pH_i_ that is known to vary by cell type, as multiple coral species express Na^+^/K^+^-ATPase, sodium bicarbonate cotransporters, and carbonic anhydrases with a high degree of tissue specificity (Barott et al., 2015b; Bertucci et al., 2011). Because acid-base homeostasis comprises diverse active and passive processes (Boron, 2004), we hypothesize cnidarian pH_i_ is governed by multiple different temperature-dependent mechanisms, a subset of which differ between cells with and without symbionts. Future studies should test the possibility that symbiont CO_2_ sequestration in the dark can still buffer pH_i_ in this species, and more generally investigate cellular and molecular determinants of thermal pH_i_ dysregulation to determine which of these are specific to cells harboring intracellular symbionts.

Both cell type and initial organismal symbiont density affected the relationship between nutritional symbiosis function and pH regulation. Non-symbiocytes from the high-symbiont-density (HD) cohort consistently had higher pH_i_ than low-symbiont-density (LD) cohort non-symbiocytes. Interestingly, this pattern was independent of host photosynthate assimilation, and is therefore unlikely to result from our hypothesized trophic differences between cohorts. Instead, this persistent difference in pH_i_ setpoint once again suggests there are cohort-level differences in *E. diaphana* cellular physiology that warrant further exploration. In symbiocytes, while there was no overall relationship between host organic carbon assimilation and pH_i_, pH_i_ specifically increased with host photosynthate assimilation only in HD anemones. This could result from temperature sensitivity in symbiont inorganic carbon uptake; that is, if the low photosynthetic rates we observed under heat stress limited symbiont CO_2_ drawdown from the cytosol, high temperatures could simultaneously lower pH_i_ in symbiocytes relative to those at ambient temperatures while also decreasing symbionts’ ability to translocate photosynthate to the host. In this scenario, non-symbiocyte pH_i_ would have decreased at high temperatures for separate reasons unrelated to symbiosis. This is consistent with our reasoning that the host and symbiont play separate roles in regulating pH_i_.

Alternatively, heat stress could have decreased symbiont productivity by disrupting host acid-base regulation of the symbiosome. If pH_i_ dysregulation in symbiotic cnidarians does not depend on bleaching, then the inability to regulate pH at high temperatures could itself contribute to symbiosis breakdown. Loss of cnidarian host control over symbiosome contents has been linked to symbiotic breakdown (Cui et al., 2019; Rädecker et al., 2018; Rädecker et al., 2021; Xiang et al., 2020), and corals use a vacuolar H^+^-ATPase to concentrate inorganic carbon in the symbiosome to promote carbon fixation by the symbiont (Barott et al., 2015a). Cnidarian host pH_i_ dysregulation, whether by ATP limitation or changes to passive buffering, could therefore 1) lead to accumulation of protons in the cytosol, decreasing symbiocyte pH_i_; and 2) contribute to the holobiont carbon limitation that is theorized to precipitate bleaching (Cunning et al., 2017; Rädecker et al., 2021; Wooldridge, 2009). This is consistent with our result that photosynthate assimilation decreased with low symbiocyte pH_i_ in the HD cohort, in which heat stress disrupted symbiotic function, but not in the putatively more heterotrophic LD cohort, which did not experience bleaching. Further research should test whether thermal acid-base dysregulation can initiate or exacerbate symbiont loss. Such a positive feedback mechanism between host and symbiont stress responses could lead to damaging synergistic effects as environmental change accelerates (Bénard et al., 2020).

## Conclusion

Here we demonstrate that thermally induced acid-base dysregulation can occur independently of bleaching in cnidarians. Heat stress appears to impose metabolic constraints on *E. diaphana* that are separate from symbiont loss, which we hypothesize disrupt their ability to regulate pH_i_. This may have negative consequences for survival of cnidarians as well as other non-endosymbiotic marine invertebrates, as the ability to reallocate ATP to pH regulatory processes is thought to be a crucial mechanism of climate change resilience (Pan et al., 2015; Sokolova et al., 2012). While endosymbiosis-specific effects on pH_i_ confirm that Symbiodiniaceae metabolism also contributes to host acid-base homeostasis, the direct effect of heat on host pH_i_ raises the intriguing possibility that thermal pH dysregulation precedes or even contributes to symbiont loss; however, additional study is necessary to validate this hypothesis. Regardless, as sea surface temperatures continue to rise (Johnson and Lyman, 2020) and marine heatwaves become more frequent and severe (Smith et al., 2023), it is crucial that we better understand how thermal stress impacts marine invertebrate cellular homeostasis. Future research on cnidarian thermotolerance should investigate specific mechanisms of ion transport disruption under heat stress. Ion transport is vital for fundamental processes including protein stability, endosymbiosis, and calcification; its disruption, while less immediately visible than bleaching, has profound negative consequences for survival (Tresguerres et al., 2014; Tresguerres et al., 2017). Understanding thermal sensitivity of ion transport and acid-base homeostasis is particularly vital as increasing atmospheric CO_2_ simultaneously acidifies and warms the oceans (Albright and Mason, 2013; Albright et al., 2016; Harvey et al., 2013). Finally, our study also demonstrates how physiological experiments in model symbiotic organisms like *E. diaphana* help address the broader question of how endosymbionts can mitigate and/or exacerbate host responses to rapid climate change, while highlighting that these effects are not uniform across laboratory populations.

## ACKNOWLEDGEMENTS

The authors would like to thank Philip A. Cleves for animal cultures and husbandry advice, Benjamin Glass and Amara Okongwu for useful discussions and respirometry assistance, and Colin Carney and the staff of the UC-Santa Cruz Stable Isotope Facility for isotope analysis assistance.

## AUTHOR CONTRIBUTIONS

Conceptualization: L.A., K.L.B.; Methodology: L.A., M.P.M., K.L.B., K.G.J., K.T.B.; Software: L.A., K.G.J.; Validation: L.A., M.P.M., K.G.J., K.T.B., K.L.B.; Formal analysis: L.A., K.G.J., M.P.M., K.T.B.; Investigation: L.A., K.G.J., M.P.M.; Resources: K.L.B.; Data curation: L.A., K.G.J., M.P.M., K.T.B.; Writing - original draft: L.A.; Writing - review & editing: K.L.B., K.T.B., K.G.J., M.P.M., L.A.; Visualization: L.A., K.G.J.; Supervision: K.L.B.; Project administration: K.L.B., L.A.; Funding acquisition: K.L.B.

## DATA AVAILABILITY

All raw data and R scripts used in data analysis are available on <u>Github</u> (https://github.com/allenwaller/Aiptasia.Heat.pHi).

